# DNA ligase Lig E increases transformation with damaged extracellular DNA

**DOI:** 10.64898/2026.03.22.713542

**Authors:** Jolyn Pan, Avi Singh, Joanna Hicks, Adele Williamson

## Abstract

Lig E is a periplasm-targeted ATP-dependent DNA ligase found in many Gram-negative bacteria including *Neisseria gonorrhoeae*. Although Lig E has been shown to have a role in biofilm formation, many Lig E-possessing bacteria are also naturally competent, suggesting a possible function in transformation with extracellular DNA. Here, we demonstrate that Lig E participates in bacterial competence by increasing transformation with nuclease-damaged extracellular DNA that contains single-stranded or cohesive breaks. We show that increased transformation with this restricted DNA is ATP-dependent, and that the ATP concentration increases in the extracellular milieu during maintenance of *N. gonorrhoeae* in liquid culture.

**Impact Statement:** Natural transformation is an important route of horizontal gene transfer that enables competent bacteria to acquire novel phenotypic traits such as antibiotic resistance or virulence factors. By demonstrating that Lig E increases transformation of *N. gonorrhoeae* with damaged resistance-encoding DNA, we provide a mechanism which competent bacteria can use to overcome nucleolytic damage sustained by environmental DNA, making this more readily available as a source of novel and potentially pathogen-enhancing genes.

## Introduction

DNA ligases seal breaks in the phosphodiester backbones of double-stranded DNA during repair, replication and recombination (1). All bacteria use an NAD^+^-dependent DNA ligase to join Okazaki fragments during DNA replication and to catalyse the final step in many DNA repair processes. However, some bacteria also encode auxiliary ATP-dependent ligases that vary in structure and function (2,3). These include Lig E, a minimal-structure DNA ligase which displays autonomous ATP-dependent DNA-joining activity despite structurally comprising only the two catalytic core domains (4). Lig E is restricted to Gram-negative proteobacteria, however, it is highly conserved and near-ubiquitous within certain taxa including Vibrionales, Campylobacterales, Neisseriales and Pasturelles (5). The *lig E* gene lacks the syntenic co-localisation with other specialised DNA repair enzymes indicating that it is not involved in dedicated stationary-phase repair pathways as other bacterial ATP-dependent DNA ligases are (5-7). Most intriguingly, all instances of *lig E* encode a 20-40 amino acid periplasmic localisation signal peptide on the N-terminus, which when deleted *in vitro*, increases both its activity and stability (4,8). This potential extra-cytoplasmic export of a DNA ligase is unexpected considering that bacterial genomic DNA is located in the cytoplasm, and suggests that Lig E could instead be acting on extracellular DNA in the periplasmic space or extracellular milieu.

Previously we reported that Lig E of *Neisseria gonorrhoeae* (Ngo-Lig E) plays a role in biofilm formation, which is consistent with both an extracellular location of this enzyme and the importance of extracellular DNA in neisserial biofilm establishment and maintenance (9,10). However we and others have observed that many Lig E-encoding bacteria are naturally competent, and are capable of taking up extracellular DNA via natural transformation and incorporating it into their genomes (7,8). Further, in both *Haemophilus influenzae* and *Vibrio cholera, lig E* is co-regulated with known competence genes, although *lig E* deletion did not cause transformation defects in either species using established assays (11,12). In the present study, we have investigated the potential contribution of Lig E activity to natural transformation in *N. gonorrhoeae*, focusing on the known biochemical activity of Lig E; its ability of join breaks in double-stranded DNA.

*N. gonorrhoeae* is an obligate human pathogen and is responsible for the second most common sexually transmitted infection in the world, gonorrhoea (13,14). *N. gonorrhoeae* is naturally competent at every stage of its growth and is able to take up pieces of extracellular DNA from the environment using its type IV pili, provided that the transforming DNA includes a specific DNA uptake sequence (DUS) (15). The high frequency of natural transformation in *N. gonorrhoeae* creates significant structural and antigenic diversity, leading to a panmictic bacterial population, and has allowed strains of this pathogen to become resistant to almost all classes of antibiotics available for disease treatment (15). The human urogenital tract where *N. gonorrhoeae* resides is a highly oxidising environment, and both reactive oxygen species produced by the host neutrophils during infection and extracellular nucleases produced by the bacterium to escape DNA-rich extracellular neutrophil traps can damage extracellular DNA, limiting its potential usefulness in transformation (16-18).

In the present work, we test the impact of Lig E deletion on DNA transformation in *N. gonorrhoeae* with both intact and nuclease-restricted DNA under non-oxidising and oxidative stress conditions. We show that Ngo-Lig E deletion has a significant impact on transformation with DNA containing single- and double-stranded (cohesive) nicks. This suggests that Ngo-Lig E could function in competence by rejoining damaged pieces of extracellular DNA, improving their suitability as a transformation substrate.

## Methods

### *Neisseria gonorrhoeae* manipulation and mutant construction

All *N. gonorrhoeae* variants used in this study are derivatives of the MS11 strain (GenBank: CP003909.1). Gonococci were grown at 37 °C with 5 % CO_2_ either on gonococcal base (GCB) agar (Difco) or in gonococcal base liquid (GCBL) (15 g/L Bacto™ Protease Peptone No. 3, 4 g/L K_2_HPO_4_, 1 g/L KH_2_PO_4_, 1 g/L NaCl), both supplemented with 1 % *Kellogg’s* supplement (22 mM glucose, 0.68 mM glutamine, 0.45 mM cocarboxylase, 1.23 mM Fe(NO_3_)_3_) (19). Piliation status was determined via morphology under a dissecting microscope at the beginning of each experiment.

To suppress pilin antigenic variation which can impact DNA uptake, a *ΔG4* derivative was generated using genomic DNA from a *ΔG4* parental strain where the G4 motif required for pilin antigenic variation has been deleted (20). The genomic DNA of a *ΔG4* strain was kindly provided by the Maier laboratory. To generate different *lig E* variants, constructs were designed to introduce insertions and/or deletions into the *ΔG4* derivative base strain by homologous recombination of flanking sequences as described previously (9,19). An *ngo-lig E* deletion mutant *(Δngo-lig E*) was created by disrupting the *ngo-lig E* gene (NGFG_RS11310) with a erythromycin resistance cassette which also provided antibiotic selection (supplementary figure S-1 A (ii) ). A complement strain was then generated by replacing the erythromycin resistance cassette in the *Δngo-lig E* mutant with a codon-optimised version of the *ngo-lig E* gene in its native site, followed by a chloramphenicol resistance cassette downstream for antibiotic selection (*ngo-lig E comp*) (supplementary figure S-1 A (iii) ). All DNA constructs were ordered as gene fragments or clonal genes from Integrated DNA Technologies (IDT) or Twist Biosciences.

Strains were generated via spot transformation as previously described (19,21). Briefly, piliated, Opa negative (Opa-) colonies were streaked through 10 ng spots of the DNA constructs. Mutants were selected on GCB agar with antibiotics (10 μg/mL erythromycin, 10 μg/mL chloramphenicol) before verification via PCR and sequencing.

### Construction of a DNA transformation reporter construct

The reporter construct for our transformation assays was generated via Golden Gate cloning (22) and consisted of a superfolder GFP reporter gene (*sfGFP*) followed by a kanamycin resistance cassette for selection (*kan*^*R*^), both under the strong constitutive gonococcal *pilE* promoter (*P*_*pilE*_). The parts were ordered as GeneBlocks from Twist Bioscience and assembled into a pTwist Amp High Copy vector backbone which contained 500 bp 5’ and 3’ homologous flanking regions to direct insertion into a hypothetical prophage region in the *N. gonorrhoeae* genome (phage protein NGFG_RS06100), and a *laczα* gene drop out for blue/white colony screening. The construct included *N. gonorrhoeae* DNA uptake sequences (5’-GCCGTCTGAA-3’) to ensure efficient uptake by the type IV pilus (23), and the plasmid backbone contained a ScaI digestion site for plasmid linearisation outside the gonococcal recombination region. Assembled reporter parts were designed with BsaI sites directing insertion into the vector backbone during cloning in the order promoter-reporter gene-antibiotic resistance gene (supplementary figure S-1 B). Parts were combined in one reaction at 18 °C overnight at a 3:1 molar ratio of each insert:backbone, in combination with BsaI-HFv2 (30 units) and T4 DNA ligase (1000 units). The reaction was terminated by incubation at 60 °C for 5 min before transformation into chemically competent DH5α *E. coli* cells. Positive clones appeared as white colonies on X-gal LB agar instead of blue, and the assembled plasmid was isolated from 5 mL of overnight liquid culture using E.Z.NA.® Endo-Free Plasmid DNA Midi Kit.

To introduce defined nuclease damage into the transforming extracellular DNA, the reporter constructs were treated with either Nb.BtsI to introduce single-stranded breaks, or NcoI to introduce cohesive double-stranded breaks. To test the impact of plasmid linearisation, constructs were treated with ScaI. All enzymes were purchased from New England Biolabs® and 10,000 ng of transformation reporter construct plasmid was digested according to the manufacturer’s instructions for each 50 μL reaction. NcoI digestion was performed at 37 °C for 2 h, followed by inactivation at 65 °C for 10 min, while Nb.BtsI and ScaI digestions were performed at 37°C for 1 h, followed by inactivation at 80°C for 20 min. The restriction digest mixtures were used directly in the transformation assays without further purification.

### Transformation assay for uptake of extracellular DNA

*N. gonorrhoeae* liquid transformation assays were performed using the established protocol of Dillard *et al*. (19). Briefly, piliated gonococci from a 24 h streak were lawned for 7.5 h. Cultures with a starting OD_550_ of 0.07 were prepared from the lawn and were left shaking overnight (30°C, 50 rpm) with sodium bicarbonate (0.042 %). 5 mL cultures with a starting OD_550_ of 0.3 were then generated and left shaking for 1.5 h (37 °C, 250 rpm), before adjusting the OD_550_ to 0.6.

The transforming DNA constructs (4000 ng) were warmed in 200 μL of GCBL media supplemented with 5.0 mM MgSO_4_ in 12-well plates for 15 min at 37 °C. Prepared gonococci were allowed to settle for 5 min and piliated cells were collected by isolating the bottom 500 μL of the inoculum. Cells adjusted to OD_550_ 0.6 (40 μL) were added to the DNA mix and incubated for 30 min at 37°C. GCBL (2 mL) was added to the transformation mix before incubation at 37°C for 2 h. The gonococci were then scrapped from the wells and serially diluted before plating onto GCB agar with 50 μg/mL kanamycin (transformant selection), or without antibiotics (total cells). The number of colonies on the agar plates were counted after 48 h to obtain colony forming unit (CFU) measurements. Transformants were also verified for sfGFP expression via fluorescence under blue light, as well as by colony PCR (with primers listed in Table S-2) to ensure proper integration into the genome. Transformation efficiencies were calculated as the CFU of transformants (on kanamycin plates) over total CFU (non-selected).

For experiments to test the effect of exogenous ATP, 1.0 mM ATP was added to the initial 200 μL of GCBL media along with the DNA of interest. For transformation after oxidative stress, gonococci were exposed to 25 mM H_2_O_2_ for 20 min. After this time cells were washed once with GCBL to remove H_2_O_2_ and resuspended in fresh media, prior to commencing the transformation protocol described above. Untreated controls were included in parallel for each treatment condition/experiment for direct comparison.

### Quantification of extracellular ATP and DNA during growth

Piliated gonococci from a 24 h streak were lawned for 7.5 h. Cultures with a starting OD_550_ of 0.07 were prepared from the lawn and were left shaking overnight (30°C, 50 rpm) with sodium bicarbonate (0.042 %). 5 mL cultures with a starting OD_550_ of 0.3 were then prepared and left shaking for 1.5 h (37°C, 250 rpm), before generation of 10 mL cultures with a starting OD_550_ of 0.08(37°C, 250 rpm). Growth was monitored hourly by measuring the OD_550_ of the culture before serially diluting and plating onto GCB agar. The number of colonies on the agar plates were counted after 48 h to obtain CFUs.

At each time point, 1 mL of the cultures were isolated for ATP and DNA quantification. These were treated with EDTA (5 mM) before heat treatment (95°C, 10 min). To isolate the cell-free extract, the samples were centrifuged at maximum speed in a benchtop centrifuge for 10 min. Extracellular ATP quantification was performed on the supernatant via a luciferase-luciferin reaction using the Invitrogen™ ATP Determination Kit according to the manufacturer’s instructions. Double-stranded and single-stranded DNA were quantified via fluorescence-based assays with the Quant-iT™ PicoGreen™ dsDNA Assay Kit and Quant-iT™ OliGreen™ ssDNA Assay Kit respectively as per the manufacturer’s instructions.

### Statistical methods

Statistical analyses were preformed using the GraphPad Prism 9.4.0 software (https://www.graphpad.com/). One-way analysis of variance (ANOVA) with Tukey’s multiple comparisons test was used to compare the different measurements and *p* values < 0.05 were deemed statistically significant.

## Results

### Lig E increases transformation with nicked extracellular DNA in *Neisseria gonorrhoeae*

To examine whether Ngo-Lig E participates in the repair of damaged extracellular DNA during competence, we transformed our *N. gonorrhoeae* variants with a DNA construct that contains single-stranded nicks in the *sfGFP* coding region (Figure 1A). This demonstrates that deletion of Ngo-Lig E significantly reduces gonococcal transformation by nicked DNA, with the *Δngo-lig E* variant having approximately 4-fold fewer transformants than when intact (non-restricted closed-circular) plasmid DNA was used (Figure 1B). However, for both wild-type and complement strains, transformation efficiencies with nicked DNA approach those with undamaged DNA, indicating that the presence of a chromosomal copy of *ngo-lig E* rescues transformation with damaged DNA. Crucially, supplementation with exogenous ATP alongside the nicked DNA significantly increases transformation of both wild-type and complement strains to levels above those observed with intact DNA. In contrast, the transformation efficiency of the *Δngo-lig E* deletion variant remains unchanged from the no-ATP condition.

**Figure 1.**
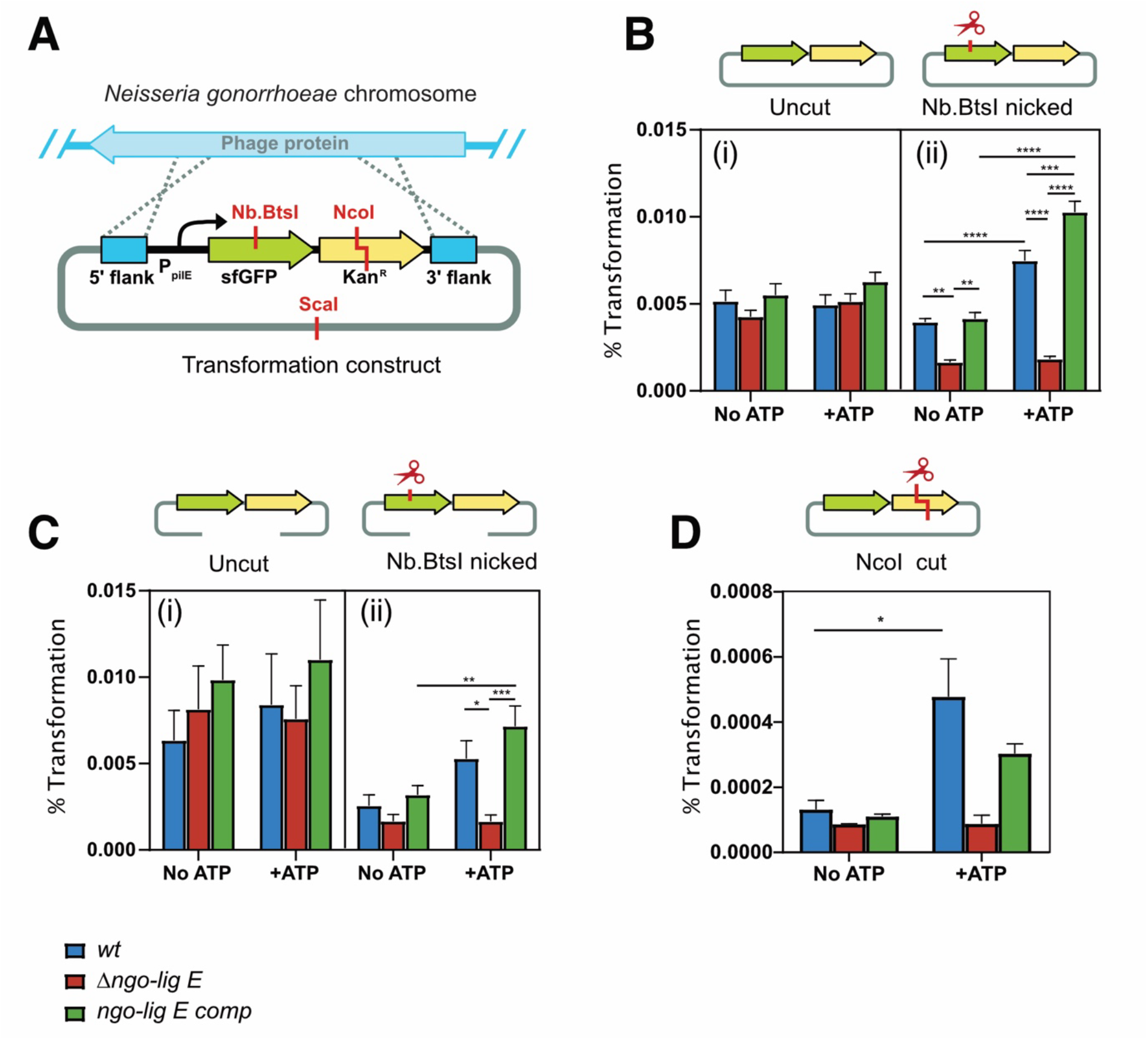
Transformation efficiencies of ΔG4 N.gonorrhoeae MS11 with constructs containing novel genes (sfGFP, kan^R^). (A) Schematic of the transforming DNA construct (not to scale) indicating general position of restriction sites used for nicking or cohesive damage. B) Transformation of circular uncut and Nb.BstI-nicked construct. (B) Transformation of linear uncut and Nb.BstI-nicked construct. (C) Transformation of NcoI cut (cohesive double-stranded overhang) construct. Transformation efficiencies were calculated as a percentage over total CFU. ‘+ATP’ refers to the supplementation of 1 mM ATP. Assays were conducted in triplicate and error bars represent the standard error of the mean with significance values given as *p ≤ 0.05; ** p≤0.01; *** p≤0.001; **** p≤0.0001. Comparisons which showed no significant differences (p > 0.05) are not indicated.

As transformation rates with intact DNA are not impacted by either differences in *ngo-lig E* status of the *N. gonorrhoeae* strain or by the addition of ATP, the increase in transformation with nicked DNA can be specifically ascribed to the ATP-dependent activity of Ngo-Lig E on the DNA damage in the wild-type and complemented strains.

We next investigated whether relaxation of the closed circular plasmid accounted for the increase in transformation efficiency by nicked DNA (Figure 1B ii) above that of intact DNA (Figure 1B i) in *ngo-lig E* -positive strains when ATP was added (Figure S-3). Linearisation of the intact extracellular DNA outside the integration region increases transformation efficiency compared with closed circular plasmid, irrespective of *ngo-lig E* status of the *N. gonorrhoeae* strain or addition of ATP (Figure 1C i). All strains are defective for transformation with nicked linearised DNA in the absence of ATP, and transformation is only restored by the addition of ATP for wild-type and *ngo-lig E* complemented strains (Figure 1B ii).

To investigate whether this enhancement of competence by Ngo-Lig E extends to repair of double-stranded damage in extracellular DNA, we repeated the transformation assay using extracellular DNA that contains a cohesive double-stranded break (Figure 1D). Transformation efficiencies for cohesive double-stranded breaks are considerably lower than with nicked DNA for all strains (∼0.0002% compared to ∼ 0.005%). However, a similar trend was observed where the transformation efficiency for wild-type and Ngo-Lig E complement strains is increased by the addition of ATP while transformation of the *Δngo-lig E* deletion strain is unchanged.

### Ngo-Lig E-related DNA uptake in *Neisseria gonorrhoeae* is not influenced by oxidative stress

Due to the oxidative nature of the human reproductive tract which may damage genomic DNA, we investigated if oxidative stress stimulates Ngo-Lig E-mediated transformation. This was done by measuring the transformation efficiencies of *N. gonorrhoeae* with intact and nicked circular DNA in the after treatment of the cells with 25 mM H_2_O_2_.

As the greatest effect of Lig E deletion was seen in the presence of ATP, we used relative ATP stimulation of transformation (% transformants with 1mM ATP / % transformants no ATP) as a proxy for Lig E activity under different oxidative stress and DNA damage conditions.

Comparisons of transformation with uncut DNA showed no significant differences between Lig E variants (wild-type, Lig E deletion or complement), or with oxidant addition (Figure 2 A). Transformation with nicked DNA showed a similar response between both oxidising and non-stressed conditions where transformation of all Lig E variants with nicked DNA was supressed in the absence of ATP, but ATP supplementation rescued Lig E-positive variants (Figure 2 B and supplementary figures S-3 and S-4). H_2_O_2_ treatment did not however change this ATP-dependent transformation recovery by Lig E positive strains. This indicates that ATP-dependent Lig E activity is not enhanced in response to oxidative stress, and neither does oxidative stress supress Lig E-mediated transformation rescue in the presence of ATP.

**Figure 2.**
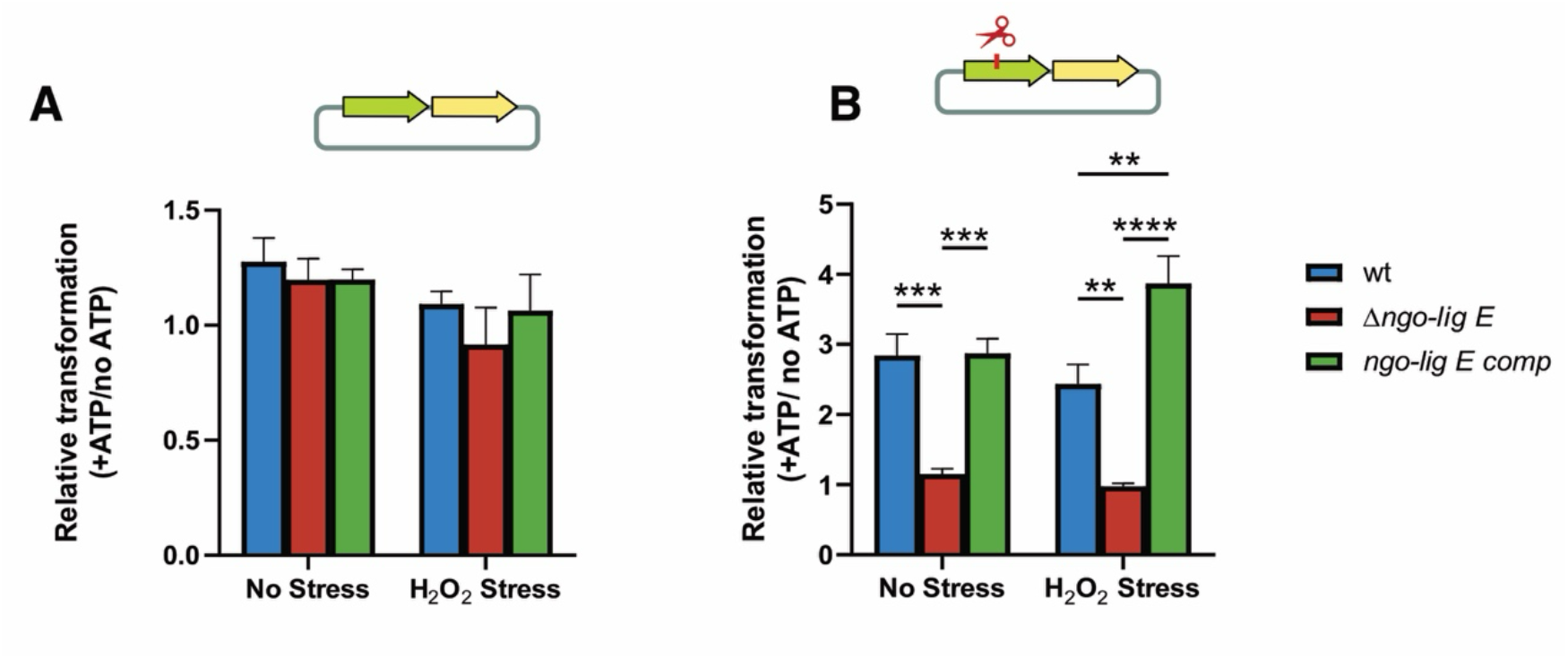
Effect of H_2_O_2_ (25 mM) on the relative transformation efficiencies of ΔG4 N. gonorrhoeae MS11 during ATP (1 mM) supplementation with constructs containing novel genes (sfGFP, kan^R^). (A) Transformation with circular uncut construct. (B) Transformation with circular Nb.Bts1-nicked construct. Relative transformation efficiencies refers to the transformation efficiencies with ATP supplementation over that of no ATP. Assays were conducted in triplicate and error bars represent the standard error of the mean with significance values given as ** p≤0.01; *** p≤0.001; **** p≤0.0001. Comparisons which showed no significant differences (p > 0.05) are not indicated.

### Substrates for Ngo-Lig E activity are available in the extracellular space

An outstanding question regarding the secretion of Ngo-Lig E is the availability of ATP in the periplasm or extracellular environment as a cofactor for ligation. To investigate this, we quantified ATP in supernatant of *N. gonorrhoeae* during normal growth in liquid media using luciferase assay (Figure 3 A, and supplementary figure S-5). This reveals an increase in ATP over time, representing a 10-fold increase in concentration by the end of the growth curve. The amount of ATP normalised to cell density was generally consistent during growth for all three *N. gonorrhoeae* mutants (S-5), suggesting that there is a constant source of extracellular ATP available for Ngo-Lig E activity in *N. gonorrhoeae*.

**Figure 3.**
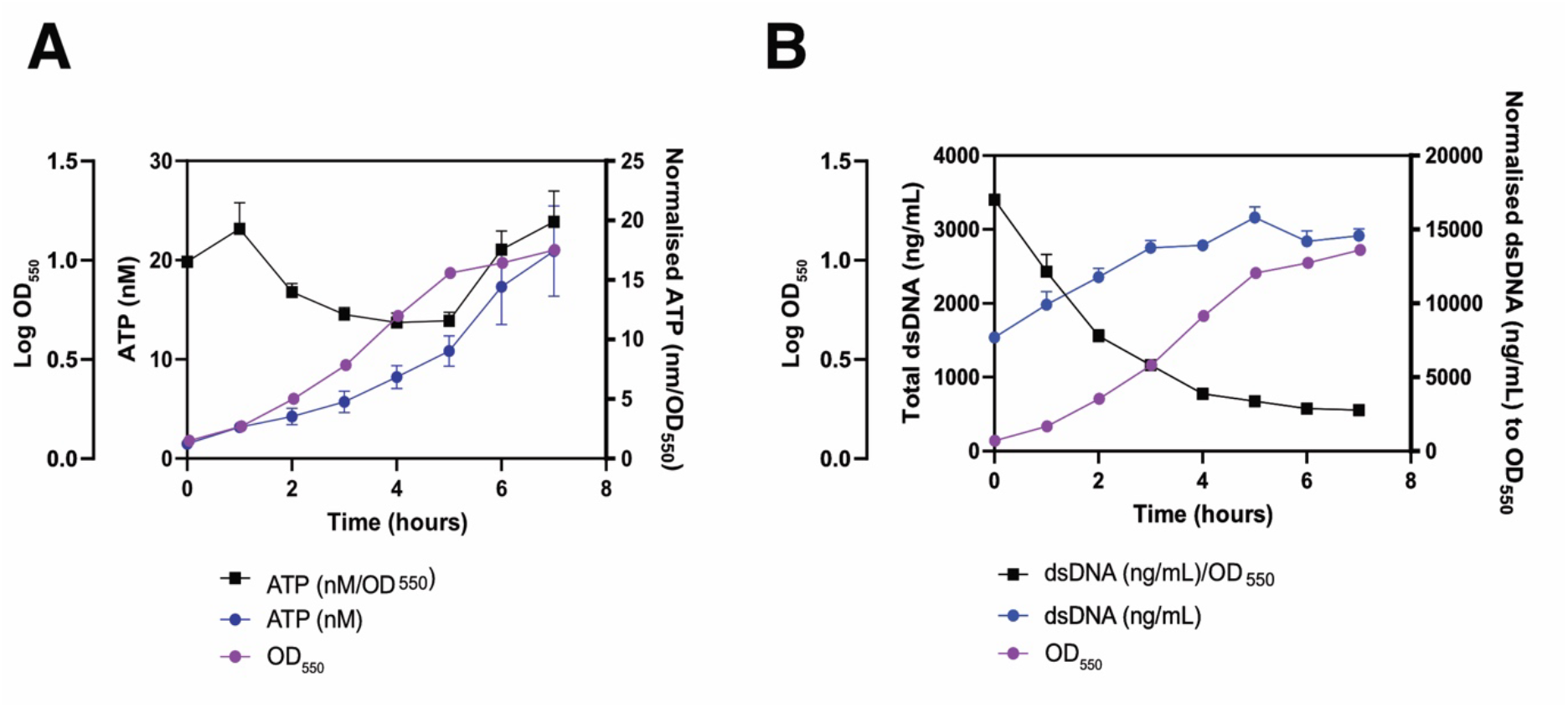
Quantification of potential Lig E substrates available in the extracellular milieu of wt ΔG4 N. gonorrhoeae during growth (A) Extracellular ATP (B) Extracellular double-stranded DNA (dsDNA). Black points are values normalised to OD at each time point; Blue points are raw values; Magenta points are OD_550_ at each time. All samples were analysed in triplicate and error bars represent the standard error of the mean Comparisons of all three lig E strains can be found in Figure S-4 to S-6

We also sought to quantify the availability of double-stranded DNA in the extracellular milieu during gonococcal growth, which could act as a substrate for transformation and potentially for Lig E activity. Using a PicoGreen dsDNA dye assay, we show extracellular double-stranded DNA increases during growth, and peaks during the exponential phase at around 3 hours before plateauing (Figure 3 B). Normalisation of extracellular DNA to cell density shows the maximum relative concentration occurs during early growth phase then decreases with time (Figure 3 B). No major differences were observed between the three strains (supplementary figures S-5 and S-6).

## Discussion

### The likely role of Ngo-Lig E on DNA uptake during natural transformation in *Neisseria gonorrhoeae*

Despite being one of the first ATP-dependent DNA ligases identified in bacterial genomes (8,24), the biological function of Lig E has remained something of an enigma. Lig E, (confoundingly annotated as Lig A in earlier literature (24,25)) was initially implicated in bacterial competence on the basis of its genetic co-localisation with competence regulatory elements (25) and predicted N-terminal signal peptide (8,26). However, *lig E* deletion in *H. influenzae* and *V. cholerae* showed no impact on either DNA uptake or transformation efficiency (11,12) and no mechanism was provided for the involvement of DNA ligase activity in competence. Here, we demonstrate that in *N. gonorrhoeae* Lig E enhances transformation specifically of extracellular DNA that contains single strand or cohesive breaks, and that *lig E* deletion significantly decreases transformation by this damaged DNA. Consistent with initial reports (11,12), *lig E* deletion has no significant impact on transformation outcomes when intact extracellular DNA is used, indicating that Lig E is not required for competence itself, but rather functions in mitigating breaks present in the transforming DNA, presumably by ATP-dependent ligation.

DNA joining by Ngo-Lig E and all other Lig E variants studied to date requires ATP as the adenylate-donating cofactor (4,8,24,27). While the bacterial periplasm was historically regarded as being devoid of ATP, more recent studies have reported nanomolar quantities in the extracellular milieu (28), and have documented active secretion of ATP by pathogenic bacteria during infection (29-31). Here we show that in *N. gonorrhoeae*, supernatant ATP increases up to a final concentration of 25 nM during liquid cultivation, averaging 15 nM per OD_550_ during growth. The detection of extracellular ATP in *N. gonorrhoeae* is in agreement with previous reports that a variety of bacteria release extracellular ATP in a growth-phase regulated manner, reaching up to 14 nM in *E. coli* (32), and ranging from 6 to 9.8 pM ATP per CFU in other Gram negative species (33). We suggest that the autolytic nature of *N. gonorrhoeae* may contribute to this pool of extracellular ATP (34) which could then diffuse across the outer membrane through porin channels into the periplasm as has been observed in *E. coli* and would then be available as a substrate for Lig E (28). Although our quantified extracellular ATP concentration is lower than the K_m_ of recombinant Lig E for ATP (0.2 μM for *H. influenzae* and 3.8 μM for *Psychromonas sp*. (4,24), and less than the ATP concentration used for supplementation experiments, it is plausible that more extracellular ATP is available in the native biological environment of *N. gonorrhoeae* during infection potentially originating from neighbouring bacteria as well as the human host cells. *E. coli*, for example has been shown to stimulate host cell release of ATP by 6-7 fold during colonisation (28).

The greatest impact of Ngo-Lig E on transformation is with singly-nicked extracellular DNA, however we also observed greater transformation of Lig E-encoding strains with DNA containing double-stranded breaks, relative to the *ngo-lig E* deletion strain. This is consistent with *in vitro* studies of recombinant Ngo-Lig E, that demonstrated singly-nicked DNA is ligated with ∼35% more efficiency than cohesive double-stranded breaks (9). However the present work does indicate that Lig E can meaningfully contribute to transformation of DNA with double-strand breaks in a biological context.

We also asked whether Ngo-Lig E contributes to transformation under oxidative stress. Although we previously showed no impact of Ngo-Lig E deletion in surviving H_2_O_2_ exposure (9), other groups have demonstrated that DNA repair is enhanced during periods of oxidative stress in *N. gonorrhoeae*, which arises from host immune responses during infection such as formation of neutrophil extracellular traps (16,35). Given these oxidative stressors will damage both extracellular and genomic DNA, we predicted that Ngo-Lig E activity could contribute directly by repairing damage resulting from oxidation-induced breaks (36) or indirectly by enhancing transformation with extracellular DNA for use in homology-mediated repair processes (37). Here we show that oxidative stress did not enhance the magnitude of Lig E-mediated transformation, suggesting uptake of repaired DNA is not important for mitigating genomic damage in this context.

### A proposed model for Lig E function in natural transformation

In line with our experimental findings, we propose that Lig E-encoding Gram-negative bacteria such as *N. gonorrhoeae* export this ligase across their cell membrane where it functions in extra-cytoplasmic DNA ligation of nicked transforming DNA, enhancing competence-mediated horizontal gene transfer.

The general steps of natural transformation are believed to be conserved between *N. gonorrhoeae* and other Lig E-encoding proteobacteria such as *H. influenzae* and *V. cholera*, where linear double-stranded DNA crosses the outer membrane into the periplasmic space via the concerted activity of the competence apparatus including the type IV pilus and ComEA protein (15,38). Then, prior to import across the inner cytoplasmic membrane, the transforming DNA is separated into single strands with one strand crossing into the cytoplasm for homologous recombination, while its complement is degraded into nucleotides in the periplasm (39-41).

We propose that by repairing single nicks or cohesive breaks in double-stranded transforming DNA, either in the extracellular milieu or in the periplasm, Lig E increases the length of single-strand DNA translocated across the cytoplasmic membrane, making it a better substrate for homologous recombination. This postulated biological function for Lig E would facilitate transformation with large DNA segments for horizontal gene transfer, potentially promoting competence-mediated acquisition of novel DNA encoding antibiotic resistance genes, virulence factors or other adaptive traits. In the absence of Lig E, nicked extracellular DNA would become fragmented when one strand is degraded, decreasing the size of single-stranded substrate entering the cytoplasm.

One likely source of DNA breaks are extracellular nucleases which are ubiquitous in the environment. Many Lig E encoding bacteria actively secrete extracellular nucleases themselves such as Dns and Xds from *V. cholera* and Nuc from *N. gonorrhoeae* which aid escape from DNA-rich neutrophil extracellular traps (18,42) and help remodel biofilm (43,44). Notably, expression of Dns in *V. cholera* decreases transformation and is downregulated upon competence induction, presumably to prevent degradation of transforming extracellular DNA (45). We propose that Lig E acts in opposition to these nucleases, potentially counteracting residual DNA-degradation during competence.

Consistent with our hypothesis that double-strand DNA repair by Lig E promotes uptake of larger gene regions, several Lig E-encoding bacteria are known to have acquired significant tracts of heterologous DNA by competence mechanisms (46). Further, the *lig E* gene is present in the genome of several high-profile antibiotic-resistant pathogens such as *N. gonorrhoeae* which are known to utilise natural transformation as a major mechanism of genetic exchange and diversification (5). For this reason, further defining a role for Lig E in increasing horizontal gene transfer from extracellular DNA via natural competence could have important implications in understanding antibiotic resistance acquisition among Gram-negative bacterial pathogens.

## Conclusions

In this report, we show that Ngo-Lig E is important at allowing *N. gonorrhoeae* to be transformed with nuclease-restricted DNA in the presence of ATP. This has important implications for *N. gonorrhoeae*’s ability to take up damaged sections of environmental DNA and hence its rapid acquisition of novel antibiotic resistance genes. We have also shown that there is a pool of external ATP and DNA substrates available for Ngo-Lig E ligation. Further work is required to determine the spatiotemporal organization of Lig E-mediated DNA repair and specifically whether Lig E is primarily localized to the cytoplasm, or exported further to the extracellular milieu. It is also of high interested to characterise the interplay between Lig E’s role in natural transformation and biofilm formation.

In conclusion, the results presented in this report highlight the importance of Lig E on the evolution of virulence and pathogenicity of *N. gonorrhoeae* specifically, and in many other proteobacterial pathogens more broadly. We anticipate that this will open up new understanding of the role of DNA repair in competence in bacteria, and in particular the ability of pathogens to be transformed with large DNA segments encoding virulence and resistance factors.

## Abbreviations

ANOVA: Analysis of Variance
ATP: Adenosine tri phosphate
bp: base pair
CFU: colony forming units
DUS: DNA uptake sequence
EDTA: Ethylenediaminetetraacetic acid
GCB: Gonococcal base
GCBL: Gonococcal base liquid
GFP: green fluorescent protein
PCR: polymerase chain reaction

## Acknowledgements

We would like to extend our thanks to Simone Remmel and Berenike Maier from the Maier laboratory (University of Cologne) for providing us with the gDNA of the *ΔG4* mutant used in this study.

## Author contributions

JP and AW conceived the project. JP generated *N. gonorrhoeae* mutant strains, conducted transformation assays and quantification of extracellular DNA/ATP and carried out all data analysis. AS designed, generated and tested the transformation reporter constructs. AW and JH assisted with design of experiments. JP drafted the initial manuscript; AW edited the final manuscript. All authors reviewed and approved the final version.

## Data accessibility

Supplementary raw data supporting these findings as well as sequences for assembly of transformation constructs are available from the authors upon request

## References

1. Zimmerman, S.B., Little, J.W., Oshinsky, C.K. and Gellert, M. (1967) Enzymatic joining of DNA strands: a novel reaction of diphosphopyridine nucleotide. Proceedings of the National Academy of Sciences of the United States of America, 57, 1841–1848.

2. Williamson, A., Hjerde, E. and Kahlke, T. (2016) Analysis of the distribution and evolution of the ATP-dependent DNA ligases of bacteria delineates a distinct phylogenetic group ‘Lig E’. Mol Microbiol, 99, 274–290.

3. Wilkinson, A., Day, J. and Bowater, R. (2001) Bacterial DNA ligases. Molecular Microbiology, 40, 1241–1248.

4. Williamson, A., Rothweiler, U. and Leiros, H.K. (2014) Enzyme-adenylate structure of a bacterial ATP-dependent DNA ligase with a minimized DNA-binding surface. Acta Crystallographica Section D: Biological Crystallography, 70, 3043–3056.

5. Pan, J., Lian, K., Sarre, A., Leiros, H.-K.S. and Williamson, A. (2021) Bacteriophage origin of some minimal ATP-dependent DNA ligases: a new structure from Burkholderia pseudomallei with striking similarity to Chlorella virus ligase. Scientific Reports, 11, 18693.

6. Pergolizzi, G., Wagner, G.K. and Bowater, R.P. (2016) Biochemical and structural characterisation of DNA ligases from bacteria and archaea. Biosci Rep, 36, 00391–00391.

7. Williamson, A., Hjerde, E. and Kahlke, T. (2016) Analysis of the distribution and evolution of the ATP-dependent DNA ligases of bacteria delineates a distinct phylogenetic group ‘Lig E’. Molecular Microbiology, 99, 274–290.

8. Magnet, S. and Blanchard, J.S. (2004) Mechanistic and kinetic study of the ATP-dependent DNA ligase of Neisseria meningitidis. Biochemistry, 43, 710–717.

9. Pan, J., Singh, A., Hanning, K., Hicks, J. and Williamson, A. (2024) A role for the ATP-dependent DNA ligase lig E of Neisseria gonorrhoeae in biofilm formation. BMC Microbiology, 24, 29.

10. Pan, J., Albarrak, A., Hicks, J., Williams, D. and Williamson, A. (2025) Influence of the ATP-dependent DNA ligase, Lig E, on Neisseria gonorrhoeae microcolony and biofilm formation. Biofilm, 10, 100292.

11. Sinha, S., Mell, J.C. and Redfield, R.J. (2012) Seventeen Sxy-Dependent Cyclic AMP Receptor Protein Site-Regulated Genes Are Needed for Natural Transformation in Haemophilus influenzae. J Bacteriol, 194, 5245–5254.

12. Jaskólska, M., Stutzmann, S., Stoudmann, C. and Blokesch, M. (2018) QstR-dependent regulation of natural competence and type VI secretion in Vibrio cholerae. Nucleic Acids Res, 46, 10619–10634.

13. Unemo, M. and Shafer, W.M. (2014) Antimicrobial resistance in Neisseria gonorrhoeae in the 21st century: Past, evolution, and future. Clinical Microbiology Reviews, 27, 587–613.

14. World Health Organization. (2024).

15. Hamilton, H.L. and Dillard, J.P. (2006) Natural transformation of Neisseria gonorrhoeae: from DNA donation to homologous recombination. Molecular Microbiology, 59, 376–385.

16. Stohl, E.A. and Seifert, H.S. (2006) Neisseria gonorrhoeae DNA recombination and repair enzymes protect against oxidative damage caused by hydrogen peroxide. J Bacteriol, 188, 7645–7651.

17. Gunderson, C.W. and Seifert, H.S. (2015) Neisseria gonorrhoeae elicits extracellular traps in primary neutrophil culture while suppressing the oxidative burst. mBio, 6.

18. Juneau, R.A., Stevens, J.S., Apicella, M.A. and Criss, A.K. (2015) A thermonuclease of Neisseria gonorrhoeae enhances bacterial escape from killing by neutrophil extracellular traps. J Infect Dis, 212, 316–324.

19. Dillard, J.P. (2011) Genetic manipulation of Neisseria gonorrhoeae. Curr Protoc Microbiol, Chapter 4, Unit4A.2-Unit4A.2.

20. Zöllner, R., Oldewurtel, E.R., Kouzel, N. and Maier, B. (2017) Phase and antigenic variation govern competition dynamics through positioning in bacterial colonies. Scientific Reports, 7, 12151.

21. Callaghan, M.M. and Dillard, J.P. (2019) In Christodoulides, M. (ed.), Neisseria gonorrhoeae: Methods and Protocols. Springer New York, New York, NY, pp. 143–162.

22. Engler, C., Gruetzner, R., Kandzia, R. and Marillonnet, S. (2009) Golden gate shuffling: a one-pot DNA shuffling method based on type IIs restriction enzymes. PLoS One, 4, e5553.

23. Goodman, S.D. and Scocca, J.J. (1988) Identification and arrangement of the DNA sequence recognized in specific transformation of Neisseria gonorrhoeae. Proceedings of the National Academy of Sciences of the United States of America, 85, 6982–6986.

24. Cheng, C. and Shuman, S. (1997) Characterization of an ATP-dependent DNA ligase encoded by Haemophilus influenzae. Nucleic Acids Research, 25, 1369–1374.

25. VanWagoner, T.M., Whitby, P.W., Morton, D.J., Seale, T.W. and Stull, T.L. (2004) Characterization of three new competence-regulated operons in Haemophilus influenzae. J Bacteriol, 186, 6409–6421.

26. Cheng, C.H. and Shuman, S. (1997) Characterization of an ATP-dependent DNA ligase encoded by Haemophilus influenzae. Nucleic Acids Research, 25, 1369–1374.

27. Williamson, A. and Pedersen, H. (2014) Recombinant expression and purification of an ATP-dependent DNA ligase from Aliivibrio salmonicida. Protein Expression and Purification, 97, 29–36.

28. Alvarez, C.L., Corradi, G., Lauri, N., Marginedas-Freixa, I., Leal Denis, M.F., Enrique, N., Mate, S.M., Milesi, V., Ostuni, M.A., Herlax, V. et al. (2017) Dynamic regulation of extracellular ATP in Escherichia coli. Biochem J, 474, 1395–1416.

29. Hironaka, I., Iwase, T., Sugimoto, S., Okuda, K., Tajima, A., Yanaga, K. and Mizunoe, Y. (2013) Glucose triggers ATP secretion from bacteria in a growth-phase-dependent manner. Appl Environ Microbiol, 79, 2328–2335.

30. Spari, D., Schmid, A., Sanchez-Taltavull, D., Murugan, S., Keller, K., Ennaciri, N., Salm, L., Stroka, D. and Beldi, G. (2024) Released bacterial ATP shapes local and systemic inflammation during abdominal sepsis. eLife, 13, RP96678.

31. Spari, D. and Beldi, G. (2020) Extracellular ATP as an Inter-Kingdom Signaling Molecule: Release Mechanisms by Bacteria and Its Implication on the Host. Int J Mol Sci, 21.

32. Mempin, R., Tran, H., Chen, C., Gong, H., Kim Ho, K. and Lu, S. (2013) Release of extracellular ATP by bacteria during growth. BMC Microbiology, 13, 301.

33. Ivanova, E.P., Alexeeva, Y.V., Pham, D.K., Wright, J.P. and Nicolau, D.V. (2006) ATP level variations in heterotrophic bacteria during attachment on hydrophilic and hydrophobic surfaces. Int Microbiol, 9, 37–46.

34. Hebeler, B.H. and Young, F.E. (1975) Autolysis of Neisseria gonorrhoeae. J Bacteriol, 122, 385–392.

35. Seib, K.L., Wu, H.J., Kidd, S.P., Apicella, M.A., Jennings, M.P. and McEwan, A.G. (2006) Defenses against oxidative stress in Neisseria gonorrhoeae: a system tailored for a challenging environment. Microbiology and Molecular Biology Reviews, 70, 344–361.

36. Balasubramanian, B., Pogozelski, W.K. and Tullius, T.D. (1998) DNA strand breaking by the hydroxyl radical is governed by the accessible surface areas of the hydrogen atoms of the DNA backbone. Proc Natl Acad Sci U S A, 95, 9738–9743.

37. Redfield, R.J. (1993) Genes for breakfast: the have-your-cake-and-eat-it-too of bacterial transformation. Journal of Heredity, 84, 400–404.

38. Seitz, P. and Blokesch, M. (2013) DNA-uptake machinery of naturally competent Vibrio cholerae. Proceedings of the National Academy of Sciences, 110, 17987–17992.

39. Chen, I. and Dubnau, D. (2004) DNA uptake during bacterial transformation. Nature Reviews Microbiology, 2, 241–249.

40. Chaussee, M.S. and Hill, S.A. (1998) Formation of single-stranded DNA during DNA transformation of Neisseria gonorrhoeae. J Bacteriol, 180, 5117–5122.

41. Matthey, N. and Blokesch, M. (2016) The DNA-uptake process of naturally competent Vibrio cholerae. Trends in Microbiology, 24, 98–110.

42. Seper, A., Hosseinzadeh, A., Gorkiewicz, G., Lichtenegger, S., Roier, S., Leitner, D.R., Röhm, M., Grutsch, A., Reidl, J., Urban, C.F. et al. (2013) Vibrio cholerae evades neutrophil extracellular traps by the activity of two extracellular nucleases. PLoS Pathog, 9, e1003614.

43. Steichen, C.T., Cho, C., Shao, J.Q. and Apicella, M.A. (2011) The Neisseria gonorrhoeae biofilm matrix contains DNA, and an endogenous nuclease controls its incorporation. Infect Immun, 79, 1504–1511.

44. Seper, A., Fengler, V.H.I., Roier, S., Wolinski, H., Kohlwein, S.D., Bishop, A.L., Camilli, A., Reidl, J. and Schild, S. (2011) Extracellular nucleases and extracellular DNA play important roles in Vibrio cholerae biofilm formation. Molecular Microbiology, 82, 1015–1037.

45. Blokesch, M. and Schoolnik, G.K. (2008) The extracellular nuclease Dns and its role in natural transformation of Vibrio cholerae. J Bacteriol, 190, 7232–7240.

46. Blokesch, M. (2017) In and out-contribution of natural transformation to the shuffling of large genomic regions. Curr Opin Microbiol, 38, 22–29.

