## Supplementary material for "DNA ligase Lig E increases transformation with damaged extracellular DNA": S-

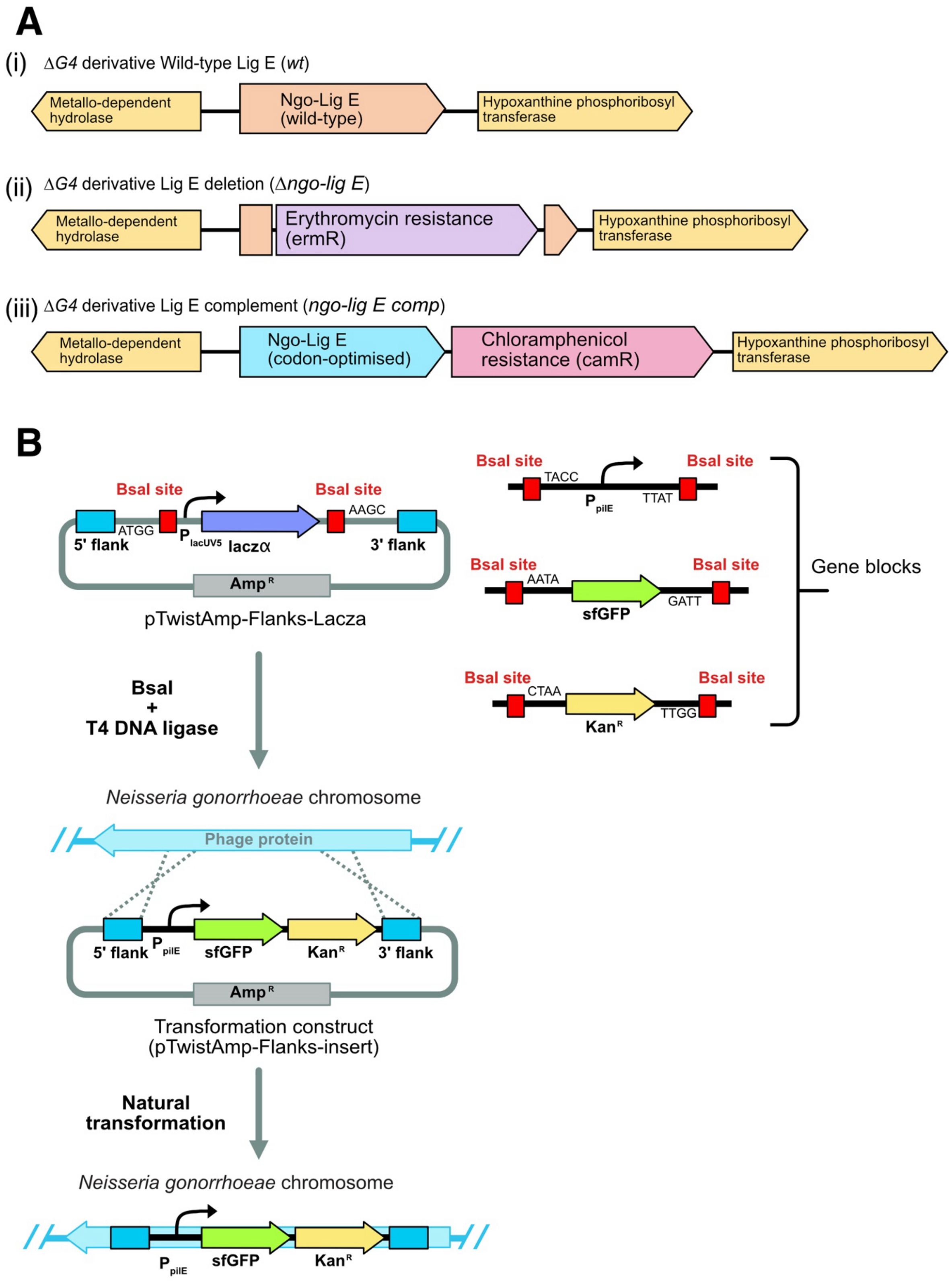


*S-1. A) Genetic context of Lig E gene for Neisseria gonorrhoeae variants used in this work. (i) Wild type with native Lig E.(ii) Lig E deletion strain. (iii) Complementation with codon-optimised Lig E under native promoter.*

*S-2. Primers used for confirmation of integration during DNA transformation assays.*

| Primer name | Sequence (5' to 3') |
|---|---|
| NS10_External_Fd | GTAACGGTTTTCAATGCC |
| NS10_External_Rev | GATATGCGCGGACATTAT |

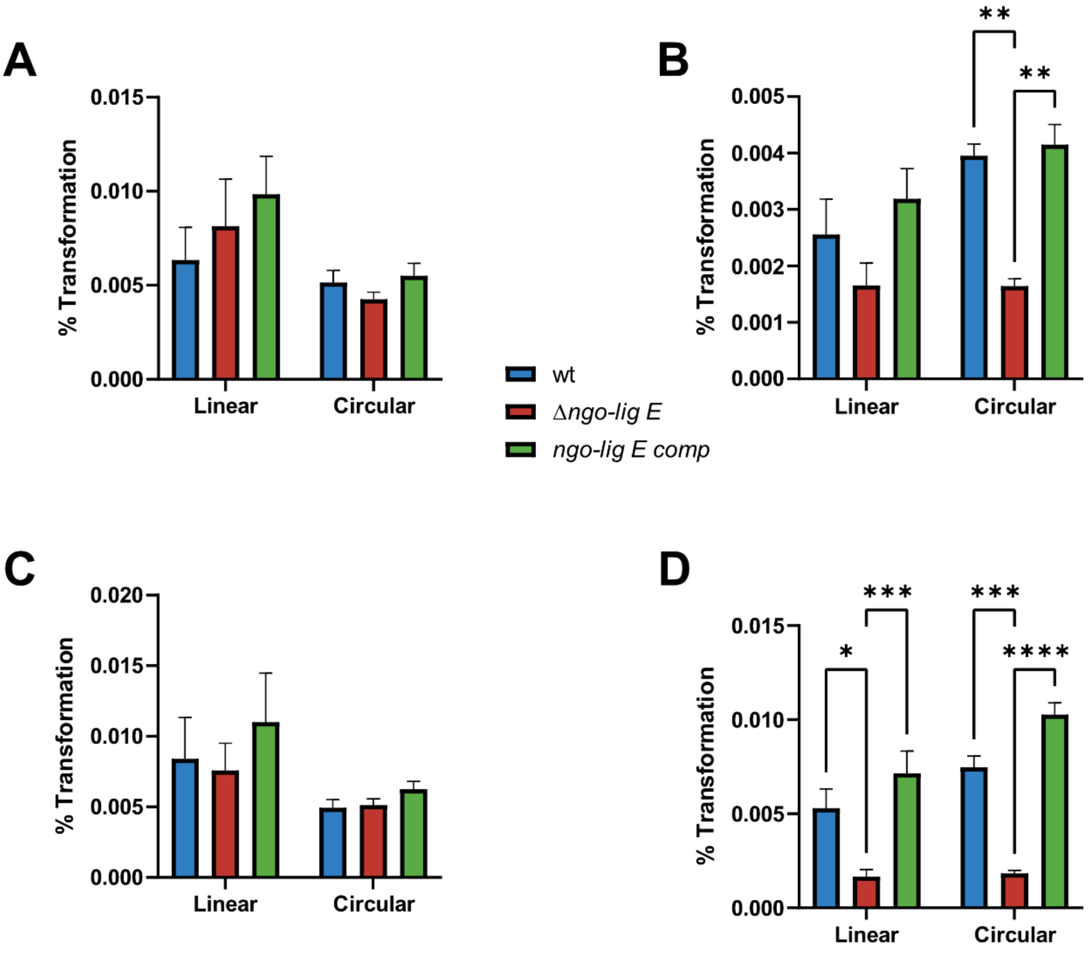


*S-3. Comparisons of transformation efficiencies of* ΔG4 N. gonorrhoeae *MS11 with either circular (non-digested) or linear (ScaI-digested) constructs containing novel genes (sfGFP, kan$^R$). A) Transformation of uncut construct. B) Transformation of Nb.BtsI-nicked construct. C) Transformation of uncut construct in the presence of 1 mM ATP. D) Transformation of Nb.BtsI-nicked construct in the presence of 1 mM ATP. Assays were conducted in triplicates and error bars represent the standard error of the mean with significance values given as *p ≤ 0.05; ** p≤0.01; *** p≤0.001; **** p≤0.0001. Comparisons which showed no significant differences (p > 0.05) are not indicated.*

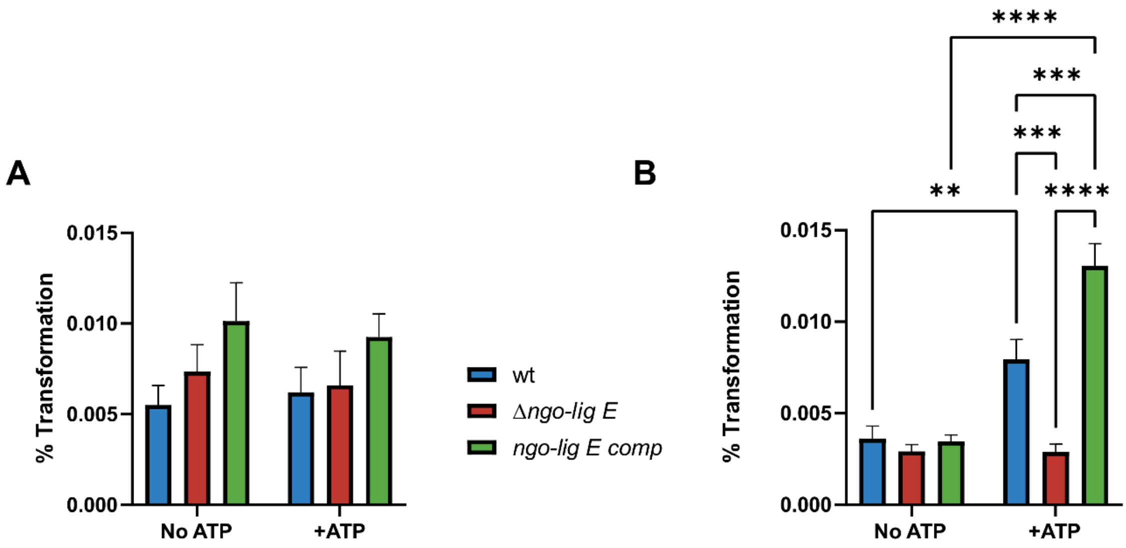


*S-4. Transformation efficiencies of $H_2O_2$ (25 mM) treated ΔG4 N. gonorrhoeae MS11 with circular constructs containing novel genes (sfGFP, $kan^R$). A) Transformation of uncut construct. B) Transformation of Nb.BtsI-nicked construct. '+ATP' refers to the supplementation of 1 mM ATP. Assays were conducted in triplicates and error bars represent the standard error of the mean with significance values given as ** p≤0.01; *** p≤0.001; **** p≤0.0001. Comparisons which showed no significant differences (p > 0.05) are not indicated.*

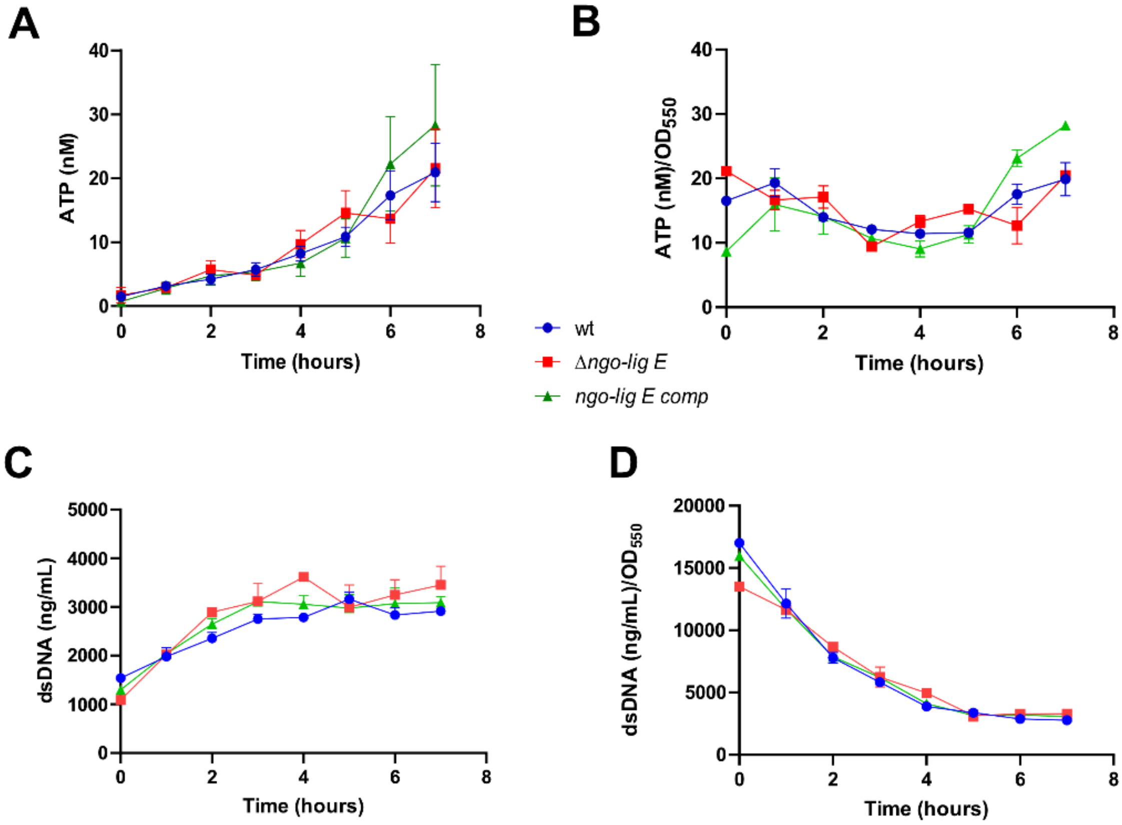

*S-5. Quantification of DNA and ATP in the extracellular milieu of lig E mutants of ΔG4 N. gonorrhoeae MS11 during growth (A) Extracellular ATP (B) Extracellular ATP normalised to $OD_{550}$ (C) Extracellular dsDNA (D) Extracellular dsDNA normalised to $OD_{550}$.*

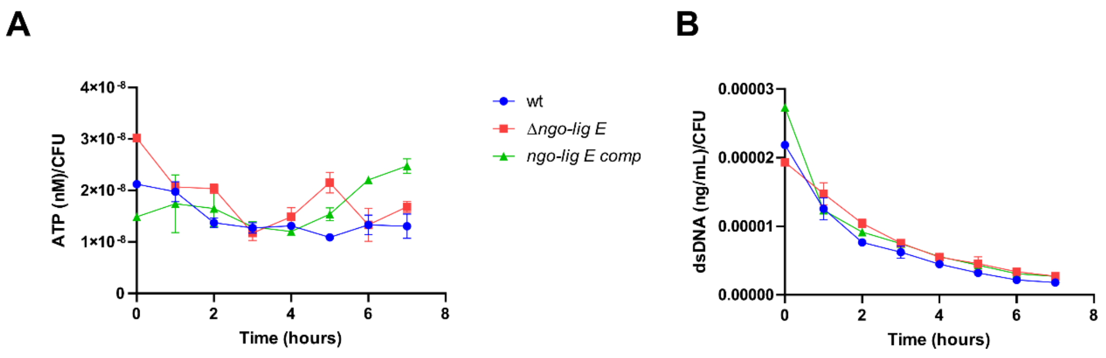


*S-6. Quantification of ATP and DNA in the extracellular milieu of* ΔG4 N. gonorrhoeae *MS11 during growth normalised to the CFU units. (A) Extracellular ATP normalised to CFU. (B) Extracellular dsDNA normalised to CFU. All N. gonorrhoeae strains were generated from the base* ΔG4 *strain.*

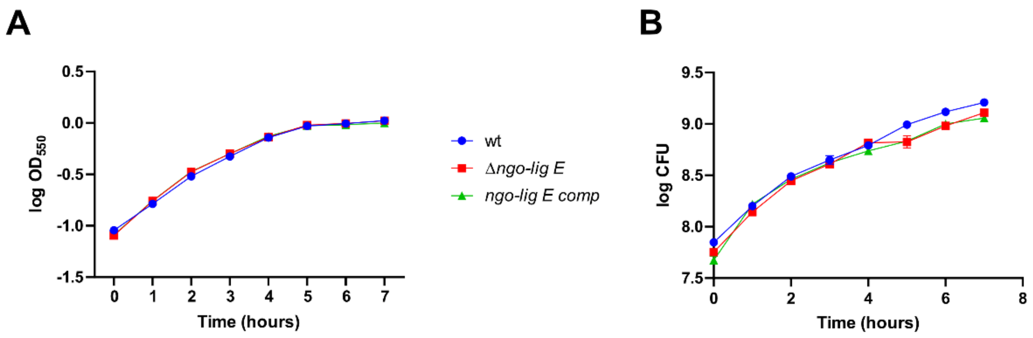


*S-7. Growth of* ΔG4 N. gonorrhoeae *MS11 varients of Lig E. (A) Log transform of $OD_{550}$. (B) Log transform of CFU counts.*
